# TCUP – An Open Access Tool to Predict Tissue of Origin and Cancer of Unknown Primary (CUP)

**DOI:** 10.1101/2025.08.08.669066

**Authors:** Ohad Landau, Eitan Rubin

## Abstract

**Introduction:** Cancer of unknown primary (CUP) remains a major diagnostic hurdle, compromising therapies that depend on accurately identifying tissue of origin. We present TCUP, an ensemble learning framework that combines Contrastive Autoencoders (CAE) and Siamese Neural Networks (SNN) with base classifiers and a meta-learning layer to classify and interpret CUP, adding biological insight through Monte-Carlo ablations.

**Methods:** Gene-expression data from TCGA (tumour), GTEx (normal), and the Genome Sciences Centre (metastatic) were imputed, log-transformed, and SMOTE-balanced. A SNN and CAE learned pairwise and reconstruction embeddings. Multiple base classifiers (e.g., SVM, Random Forest) generated meta-features, which a meta-learner combined for final prediction. Monte-Carlo ablation iterations were performed to assess gene-level importance.

**Results:** TCUP achieved 98.3 % accuracy (F1 = 98.3) across all tissues. In metastatic BRCA, COAD, and PAAD it reached 86.7 % accuracy. Ablation highlighted 79 key contributors, including established tumour suppressors NKX6-1 and SOX30 and the less-studied SYTL1. PCA confirmed clearer separation in embedded space.

**Conclusion:** TCUP delivers high tissue-of-origin accuracy and CUP assignment while providing interpretable gene importance that clarifies tissue differences and metastatic drivers. By integrating advanced embeddings with systematic ablation TCUP supplies an accessible framework to advance CUP research and, ultimately, improve clinical outcomes. TCUP is freely available at https://fohs.bgu.ac.il/rubinlab/TCUP/

## 1 Introduction

Cancer of unknown primary (CUP) is defined as “a case in which cancer cells are found in the body, but the place where the cells first started growing (origin) cannot be determined; also called carcinoma of unknown primary” [1]. Although CUP represents only 3–5 % of all metastatic malignancies [2], its clinical outcome is dismal: population-based analyses report median overall survival of 8–11 months and a one-year survival rate of roughly 25 % [3, 4]. By definition, CUPs present as advanced, heterogeneous metastases, and even exhaustive histopathologic and radiologic work-ups rarely reveal the primary site [5]. The absence of an identified origin restricts the use of site-directed therapies [6]; nevertheless, emerging precision-oncology approaches that target actionable mutations may benefit a subset of patients [7].

Conventional evaluation of CUP begins with an exhaustive clinicopathologic work-up. Extended panels of immunohistochemical (IHC) stains are used to narrow the differential diagnosis, yet large series indicate that IHC establishes a confident tissue of origin (TOO) in fewer than one-third of patients and can yield conflicting interpretations that require expert adjudication [4, 6, 8]. Cross-sectional imaging with contrast-enhanced computed tomography is routine, and ^18^F-FDG PET/CT is recommended when initial imaging is unrevealing; nevertheless, the primary lesion is visualized in only ≈ 20–30 % of cases even with PET [8, 9]. Consequently, most patients remain true CUP and receive empiric, broad-spectrum chemotherapy— typically carboplatin–paclitaxel or cisplatin–gemcitabine—as endorsed by current guidelines [8].

Molecular classifiers provide an increasingly valuable adjunct to this paradigm(see below). Numerous tissue-of-origin (TOO) classification algorithms have been developed across multiple omics platforms, leveraging the diverse molecular information contained in these data types. Genomic approaches exploit somatic-mutation signatures or copy-number profiles [10–14]; transcriptomic methods mine gene-expression patterns [15–21]; and epigenomic classifiers analyse DNA-methylation landscapes [22–24]. More recently, deep convolutional neural networks have produced promising results from cytology images [25,26], most notably, Tian et al. demonstrated that a cytology-based model significantly outperformed both junior and senior pathologists across multiple metrics when classifying CUP samples [25].

Collectively, these studies illustrate the potential of machine-learning and deep-learning techniques to tackle the disproportionate mortality burden associated with the CUP patient population [3]. However, TOO predictors are still absent from day-to-day clinical protocols. This limitation stems in part from the fact that most published models were trained and validated on large collections of well-annotated primary tumours or resected metastases rather than on true CUP biopsies, limiting their generalizability to the highly heterogeneous CUP population [6]. Only sparse data involving prospective studies demonstrating a survival benefit for CUP through algorithm guided, site-specific therapies exist [27], thus current guidelines continue to recommend empirical platinum-based doublets for the majority of patients [8]. In addition, marked variability among sequencing platforms, bioinformatic pipelines and gene-panel coverage hinders inter-laboratory reproducibility, a challenge underscored by large pan-cancer comparisons of technical heterogeneity [28, 29].

In this study, we present TCUP, a novel framework that leverages > 21 000 transcriptomic profiles from healthy tissues, primary tumors, and metastases. TCUP first learns a generalized embedded space that delineates tissue types based on gene-expression signatures; these robust representations are then fed to conventional machine-learning classifiers, whose outputs are integrated by a meta-learner to infer the tissue of origin of metastases. Monte Carlo ablation analysis subsequently identifies the relative impact of removing information at the gene-level to prediction accuracy, thereby enhancing interpretability and broadening its research and clinical utility. Lastly, a user-friendly interface facilitates straightforward deployment, promoting reproducibility and potential clinical adoption.

## 2. Methods

### 2.1 Overview

A novel pipeline was developed for identifying CUPs. Expression profile of a tumour with unknown origin is processed by sequential models. A Siamese neural network learns latent representations in which samples from identical tissue types cluster. In parallel, a contrastive autoencoder generates an independent embedding set by reconstructing gene-expression signals while jointly minimizing reconstruction and contrastive losses, yielding additional tissue-discriminative features. The two embedding spaces are concatenated to form a unified low-dimensional feature matrix that is supplied to an ensemble of classifiers trained to predict tissue of origin (TOO). Class probabilities from these classifiers are integrated by a meta-learner to produce the final TOO assignment

### 2.2 Data

Datasets were collected from BCGSC [29] and TCGA [30], obtained through the cBioportal [31], accessed February 2025 and Genotype-Tissue Expression (GTEx) [32], accessed February 2025. Our final dataset comprised of 4,380 tumor samples from 15 different cancer types, balanced to ensure equal representation per type. Advanced metastatic tissue transcriptomics were obtained from a Pan-cancer analysis of advanced patient tumors [29] and filtered to match TCGA cancer types and maintained only samples where the metastatic lesion site is different from the primary – we defined as CUPs. This resulted in 14 different metastatic classes, 313 total samples, of which 4 classes had sufficient cases (n>5) to be incorporated into a test split—BRCA (40), COAD (35), LUAD (20), and PAAD (25). Finally, we obtained 16,245 healthy tissue transcriptomics from GTEx of 26 different tissues. In total, 21,321 samples and 45 labels were used, stratified, for the various training, validation and testing splits, as shown in Table 1. All transcriptomics obtained were normalized to transcripts per million, then log transformed. We employed LASSO-based feature selection (2548 genes) [35] and used a standard scaler fitted on the training set and applied to the validation and test splits.

**Table 1:** Demographic and clinical data. Distribution of samples and their demographics across the difference cancer types (classes) and splits.

| Class | Total (n=) | Median Dx Age (yrs) | % Female | Median OS (months) | % Dead |
| --- | --- | --- | --- | --- | --- |
| BLCA | 299 | 69.0 ± 10.8 | 27.4 | 17.7 ± 28.1 | 45.2 |
| BRCA | 300 | 60.0 ± 13.7 | 99.7 | 28.2 ± 41.4 | 15.0 |
| CESC | 287 | 46.0 ± 13.9 | 100.0 | 20.9 ± 37.0 | 23.0 |
| COAD | 299 | 68.0 ± 13.1 | 48.1 | 20.2 ± 23.4 | 21.6 |
| HNSC | 299 | 61.0 ± 12.1 | 28.1 | 21.8 ± 29.6 | 44.8 |
| KIRC | 299 | 61.0 ± 12.2 | 39.5 | 43.2 ± 32.3 | 33.8 |
| LGG | 299 | 42.0 ± 13.0 | 44.8 | 24.8 ± 33.4 | 23.7 |
| LIHC | 299 | 61.0 ± 13.4 | 35.1 | 19.6 ± 23.9 | 35.5 |
| LUAD | 299 | 66.0 ± 9.9 | 54.5 | 21.6 ± 27.4 | 34.8 |
| PAAD | 179 | 65.0 ± 10.9 | 44.7 | 15.3 ± 15.6 | 52.0 |
| PRAD | 299 | 62.0 ± 6.5 | 0.0 | 29.2 ± 26.5 | 1.3 |
| SKCM | 276 | 57.0 ± 16.0 | 38.0 | 38.0 ± 65.0 | 46.5 |
| STAD | 299 | 67.5 ± 10.9 | 36.8 | 14.4 ± 16.8 | 40.8 |
| THCA | 293 | 47.0 ± 16.3 | 74.1 | 31.3 ± 32.9 | 3.8 |
| UCEC | 299 | 64.0 ± 11.2 | 100.0 | 29.7 ± 30.2 | 17.4 |
| Metastatic | 313 | 57.0 ± 11.2 | 70.0 | 26.2 ± 29.1 | 76.4 |
| Train | 2736 | 61.0 ± 14.4 | 51.7 | 23.4 ± 34.1 | 31.0 |
| Validation | 919 | 60.0 ± 14.1 | 51.0 | 23.2 ± 34.3 | 31.4 |
| Test | 983 | 60.0 ± 14.1 | 51.3 | 24.0 ± 34.1 | 34.4 |
| GTEX | 16196 | NA | NA | Not relevant | Not relevant |

**Note:** The 16196 GTEx healthy-tissue samples are distributed across 26 tissue labels, while the 313 metastatic samples are distributed across 8 metastatic tissue labels.

### 2.3 Siamese Neural Networks and Contrastive Autoencoder

We trained both a Siamese Neural Network (SNN)[33] and a Contrastive Autoencoder (CAE) [34] using a 60 - 20 - - 20; train–validation–test split (Table 1). To improve efficiency, we used LASSO to select 2,548 genes [35]. For the SNN and CAE, randomly generated pairs of samples were labeled positive when they belonged to the same tissue or cancer type and negative when they did not. To limit overfitting, each sample appeared in no more than five positive and five negative pairs. Because metastatic specimens were under-represented, we applied SMOTE oversampling solely to the training set samples of those classes and further compensated by permitting up to thirty pairings per metastatic sample. The SNN was optimized with a binary cross-entropy loss computed on the latent-space distance of each pair. By contrast, the CAE was trained with a dual objective that combined a margin-based contrastive loss with a reconstruction loss (mean-squared error) using Pythons TensorFlow library. Jointly optimizing these objectives, together with shared embeddings, yielded a more informative latent space (Supp. S1). Metastatic labels represented by fewer than five samples were included only in the pairing logic of the training and validation sets—never in classification—to enrich the latent representation. The SNN was trained for 40 epochs with early stopping, whereas the CAE training extended to 60 epochs. To address the heightened heterogeneity of metastatic cases and other outliers, we employed hard-example mining (HEM) in two targeted passes, followed by a final reweighting stage:

● **Initial pass (10 epochs)**: a straightforward 10-epoch run over the entire training set.
● **Uncertainty HEM (10 epochs)**: the 15 % of pairs whose predicted probabilities lay closest to 0.5 (i.e., most uncertain) were isolated, and the model was retrained solely on these examples.
● **Overconfidence HEM (10 epochs)**: a different 15 %—pairs misclassified despite high predicted confidence—were selected for exclusive retraining.
● **Final reweighting (10–30 epochs)**: training resumed on the full dataset, allowing the model to converge while balancing average and hard examples.

The learning curves demonstrate that hard example mining effectively boosted performance, with the sharpest gain achieved during the CAE’s uncertainty pass (Fig. 1E).

**Fig. 1:**
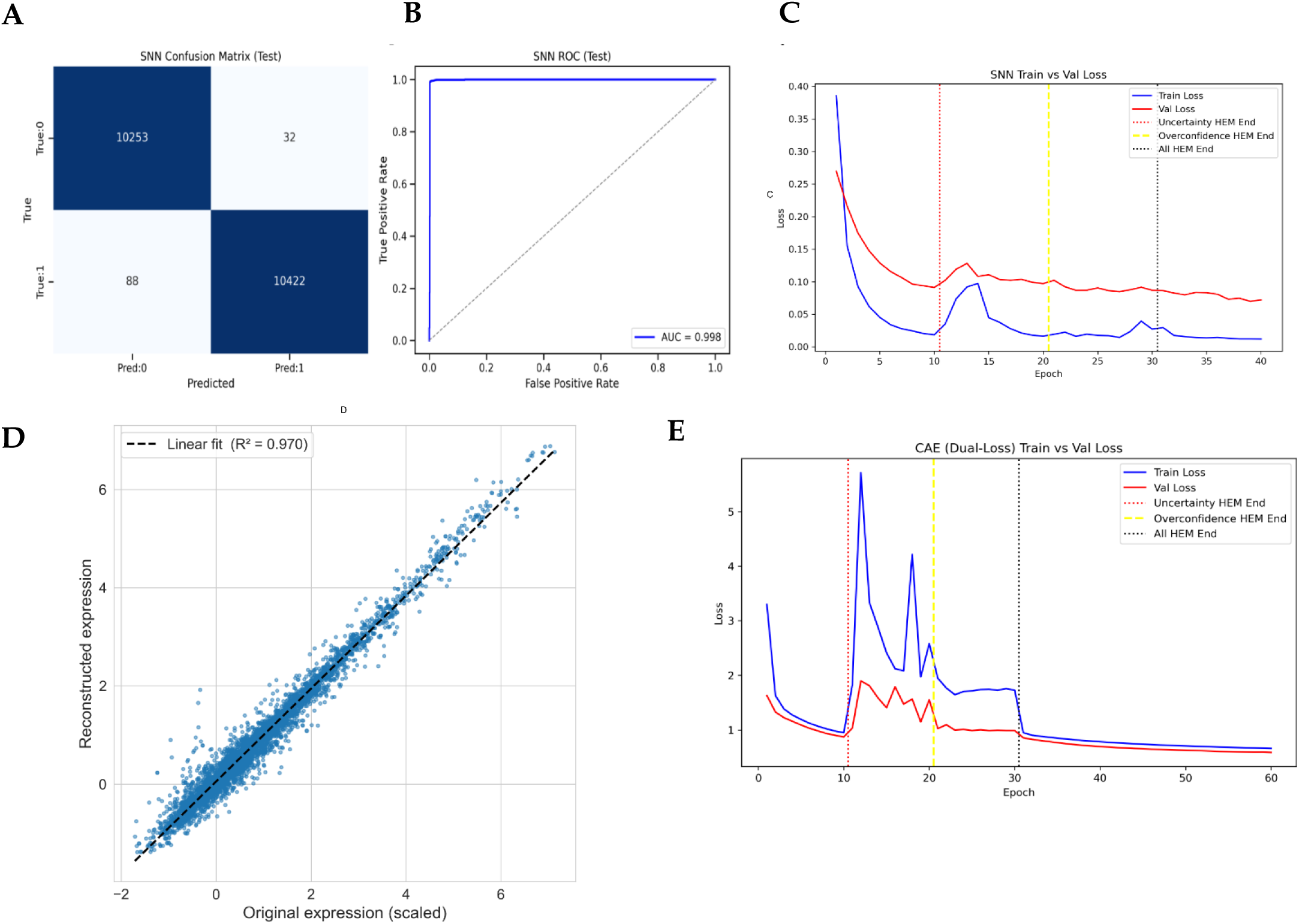
Training and Evaluation of the Siamese Neural Network (SNN) and Contrastive Autoencoder (CAE). A – Confusion matrix showing SNN predictions of similar or dissimilar for a given pair. B – Receiver operating characteristic (ROC) curve for the SNN; AUC = 0.998. C – SNN loss curve, highlighting Hard Example Mining (HEM) stages. D – Scatter plot of Original and Reconstructed expression, R^2^=0.970. E – CAE loss demonstrating extended convergence through HEM phases.

### 2.4 Meta Learner

After training the SNN and CAE, the data was split once into an 80 % development set and a 20 % independent test set. Tuning of hyperparameters was carried for the five base classifiers—logistic regression (LR), k-nearest neighbours (KNN), random forest (RF), support-vector machine (SVM), and XGBoost (XGB) – on the development set (70-30 split) using Pythons Sklearn library. Predicted class labels and their probabilities were concatenated to create a meta-feature matrix. This matrix was then used to tune the parameters of the meta-learner (60-20-20 splits, development set). Finally, the untouched 20 % test set was embedded by the SNN and CAE, classified by the five base models, and the outputs combined by the meta-learner, which outperformed every individual classifier (Supplementary Fig. S2).

### 2.5 Monte Carlo Simulations

To quantify feature importance, we carried out a Monte Carlo single-gene ablation study. The 2548 genes (see methods 2.3) were individually perturbed by adding Gaussian noise (μ = 0, σ = 20) to its standardized expression profile; the full classification pipeline was then re-executed and the consequent change in accuracy recorded. Repeating all 2,548 ablations ten times averaged out stochastic variation and yielded a robust estimate of each gene’s contribution to tissue-of-origin prediction, also analyzed separately for GTEx, TCGA, and metastatic cohorts and for class specific effects. Genes were subsequently ranked by the mean accuracy drop. Then, the 128 most influential genes (ranked-based p-value; p<0.05) were ablated in combination as well as the 128 least influential genes or ten random 128-gene cohorts. We then iteratively pruned the ranked list—from least to most influential—to identify the optimal subset whose removal produced a maximal and significant (p<0.05; McNemar’s χ² test) accuracy decline.

### 2.6 Selecting most important subset of genes

Monte-Carlo gene ablation isolated label-specific markers. The two most accurate classes—BRCA (98.3 %) and metastatic BRCA (97.5 %)—were analyzed. For each class, the accuracy drop produced by ablating each of >2,500 gene alone was ranked by p-value; the 128 top-ranking genes (p < 0.05 for all) formed the initial candidate lists. Retaining only genes significant for metastatic BRCA but not for BRCA yielded 110 putative metastatic genes. The entire 110-gene panel was then ablated collectively and iteratively pruned by reinstating genes in ascending order of individual effect until the subset that maximized class contrast was reached: a significant accuracy loss for metastatic BRCA (p < 0.05) with no significant loss for BRCA (p > 0.05). This optimized subset was designated the final differentiating signature and provided the foundation for a focused literature review of each gene’s function and cancer associations.

### 2.7 User Interface

To enable external deployment, the complete TCUP pipeline was implemented as a Dash web application (https://fohs.bgu.ac.il/rubinlab/TCUP/) and hosted on the Render platform. Two reference expression panels support on-line imputation of TOO: a Cancer cohort (TCGA + metastatic samples) and a Healthy cohort (GTEx). For every gene, the log-transformed cohort medians were pre-computed and stored as an imputed value. Uploaded transcriptomes undergo the same log-transformation and global standard-scaling parameters defined during model development then trimmed to match selected features; any missing gene is replaced by the corresponding cohort median. The resulting feature vector is encoded by the pretrained SNN and CAE, passed through the five base classifiers, and scored by the meta-learner to yield tissue-of-origin probabilities. The application reports the predicted class and a ranked list of the twenty genes whose single-gene ablation produced the greatest accuracy drop in Monte-Carlo analysis, together with each gene’s standard deviation (σ) z-score (standard deviation distribution per gene per cohort) relative to the selected reference cohort.

## 3. Results

### 3.1 Building TCUP

Transcriptomic profiles from GTEx, TCGA, and BGCSC were processed as outlined in Methods 2.1, after which LASSO regularization reduced the original feature set to 2,538 genes (Fig 2I). Pair generation ensued to create a dataset of similar and dissimilar samples, with similarity defined by the tissue source (Fig 2II). Then, the genes were embedded by two independent networks — a Siamese Neural Network (SNN) and a Contrastive Autoencoder (CAE). The reduced SNN and CAE embeddings served as input to five base classifiers (Fig 2IV), whose class-probability outputs were merged and fed to a meta-learner (Methods 2.3); the schematic of this architecture is provided in Fig. 2. Evaluation on the 20 % held-out test set confirmed good generalization, with the SNN alone achieving clear separation of similar versus dissimilar tissues (Fig. 1A-C). During training, each Hard-Example Mining (HEM) phase (10-30 epochs) produced a transient rise in loss, reflecting the increased difficulty of the curated mini-batches, yet every HEM cycle concluded below its initial loss at the last epoch, documenting consistent performance gains (Fig. 1C,E). The narrow and stable gap between training and validation losses further indicated negligible overfitting. In parallel, the CAE was able to reconstruct gene-level expression profiles with high fidelity (R^2^ = 0.970), underscoring the biological coherence of its latent space (Fig. 1D).

**Fig. 2:**
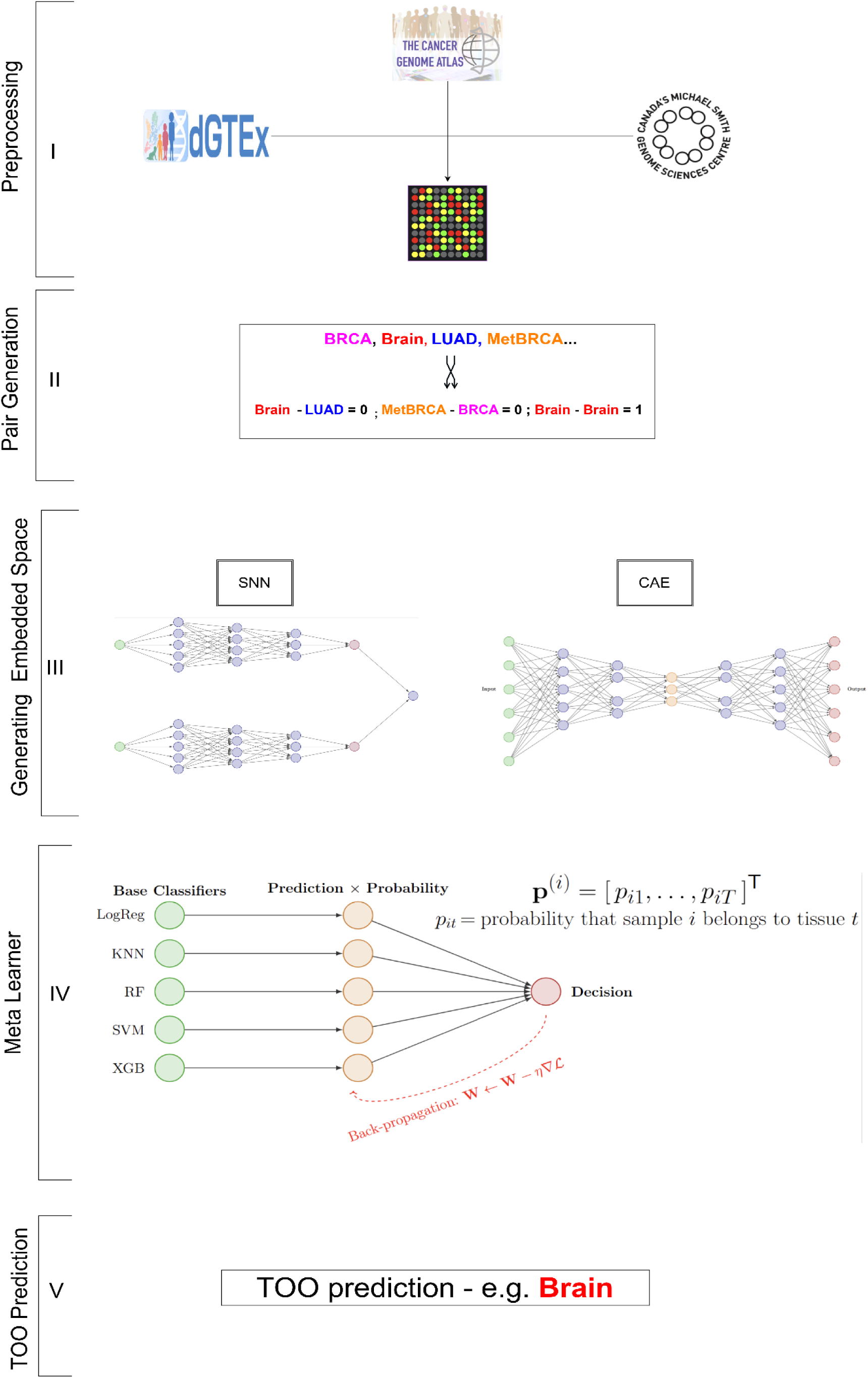
Overview of the TCUP classification pipeline. I – Preprocessing, incorporating the various datasets. II – Pair Creation - positive pairs represent samples of the same tissue; negative pairs represent any combination of different tissues. III – SNN and CAE trained on recently created pairs to generate embeddings. IV - Meta-learner incorporates base learner’s predictions on embedded space and fine tunes predictions. V – TOO classification based on highest probability class.

### 3.2 Evaluating TCUP

Using the architecture outlined in Section 3.1, we evaluated performance on the independent 20 % test partition. Across 45 tissue, tumor, and metastatic labels, the ensemble achieved an overall accuracy of 98.3 % and an F1-score of 98.3 %, comparable with recently published (>2022) algorithms (Table 2). A salient error emerged in one non-metastatic class: healthy breast tissue, where accuracy declined to 92 % and misclassifications were predominantly assigned to healthy adipose tissue (Fig. 3A; Supp. Fig. S3). For metastatic samples, the mean accuracy was 86.7 % (Fig. 3B), increasing with larger class sizes and reaching 97.5 % for metastatic BRCA. Receiver-operating-characteristic (ROC) analyses corroborated these findings, yielding an area under the curve of 1.00 for the complete 45-class model and 0.96 when restricted to metastatic labels (Fig. 3C). Finally, principal-component projections of raw expression profiles versus latent embeddings revealed markedly reduced cluster overlaps in both two- and three-dimensional space, indicative of superior class separability within the learned feature space (Fig. 3E).

**Fig. 3.**
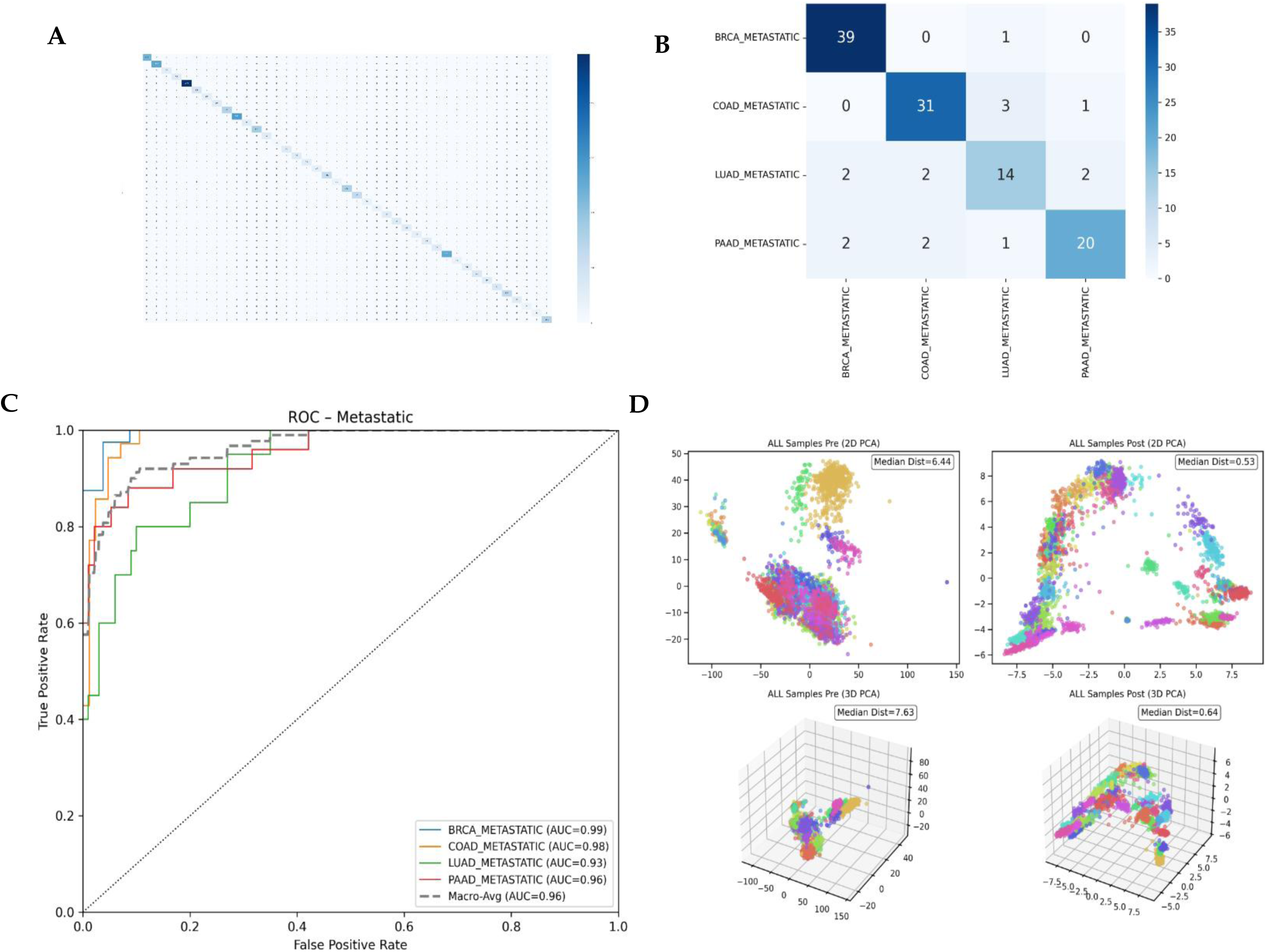
– TCUP evaluation. A – 45 predicted vs. true confusion matrix; the heat signature corresponds to accuracy per label (‘Zoomed-in’ at Supp S3-S7). B –Metastatic-specific confusion matrix. C - Metastatic specific ROC curves. D – 2D and 3D PCA, pre and post embedded space, markedly increasing clusters quality and tissue separation (Median distortion – 2D – 6.44 vs 0.53; 3D – 7.63 vs 0.64), colors representing the 45 different labels.

**Table 2.** Performance of TCUP vs Published algorithms.

| Model / Dataset | Accuracy (%) | Top-2 Acc. (%) | Top-3 Acc. (%) | F1 Score (%) | Recall (%) | ROC-AUC |
| --- | --- | --- | --- | --- | --- | --- |
| TCUP – Full* / ** | 98.3 | 99.2 | 99.5 | 98.3 | 98.3 | 1.000 |
| TCUP – TCGA* | 96.2 | 97.8 | 98.2 | 96.2 | 96.2 | 0.997 |
| TCUP – GTEx | 99.3 | 99.8 | 99.9 | 99.3 | 99.3 | 1.000 |
| TCUP – Metastatic (CUP)** | 86.7 | 93.3 | 97.5 | 86.4 | 86.7 | 0.965 |
| He *et al.* (RNA)<sup>[19]</sup> | 97.5*/82.7**–91.1** | — | — | — | — | — |
| Rydzewski *et al.* (RNA)<sup>[20]</sup> | 92.1*/86.8** | — | — | — | — | — |
| Divate *et al.* (RNA)<sup>[21]</sup> | 97.33*/88** | — | — | 97.33* | 97.33* | — |
| Nguyen *et al.* (DNA)<sup>[13]</sup> | 89** | 94** | — | — | 90** | — |
| Brlek *et al.* (DNA)<sup>[14]</sup> | 81* | 91* | — | 81* | — | 0.970* |
| Lu *et al.* (Image)<sup>[25]</sup> | 83** | — | 96** | — | — | — |
| Tian F *et al.* (Image)<sup>[26]</sup> | 82.6** | — | 98.9** | — | — | 0.969** |
| Duckett *et al.* (Epigenetics)<sup>[23]</sup> | 97.2*/91.5** | — | — | — | — | 0.952*/0.915** |
| Chen *et al.* (Epigenetics)<sup>[24]</sup> | 97.2* | — | — | — | — | — |

\* Tissue of Origin (TOO): Model predicts original tissue type from primary or non-metastatic tumor samples.
\*\* CUP model accuracy when inferring origin of metastatic samples.
- While the metrics reported in this table represent core performance values, they do not fully capture the broader scope of each work. Several works expanded beyond classification, incorporating clinical tools, comparisons with pathologists, evaluations on CUP-specific data, demonstrations of improved clinical outcomes, and more. The values presented reflect only the raw comparative metrics of the selected algorithms.

### 3.3 Monte Carlo Ablation

To interpret the TCUP model, gene ablations were performed. For each of the genes, Gaussian noise was injected, and the modified inputs were passed through the full TCUP pipeline. The drop in overall accuracy indicated each gene’s significance for tissue-of-origin classification, averaging the effect of 10 ablation repeats per gene to reduce stochastic variation. Next, an optimal, minimal gene subset was identified whose ablation produced the greatest performance drop across all tissue types. Briefly, this was achieved starting with the 128 most influential genes (significantly effecting accuracy; p < 0.05). All were ablated, and the resulting accuracy was recorded. We then repeated the procedure while iteratively reinstating one gene at a time. At each step, the observed accuracy loss was contrasted against that produced by ablating (i) the bottom-n genes and (ii) ten random n-gene cohorts of equal size (Methods 2.4). This analysis pinpointed 79 genes whose collective removal impaired performance substantially more than all controls (p < 0.0001; Fig. 4A). An additional metastatic-specific evaluation of TCUP performance identified a 40-genes set whose joint ablation significantly reduced accuracy (p<0.01, McNemar’s χ² test test) for metastatic compared to the random or bottom-ranked controls (p < 0.01, McNemar’s χ² test; See Supp. Fig. S8 for more details). The twenty genes with the greatest individual impact on accuracy are listed in Fig. 4B, and for the most influential 4 (shown in red in Fig. 4b) the normalized expression patterns across tissues are visualized in Fig. 4C. Although these box plots reveal pronounced tissue-level separation, the variability among individual points (samples) highlights the multivariate nature of the task: no single gene transcript is sufficient to resolve tissue of origin in isolation.

**Fig. 4:**
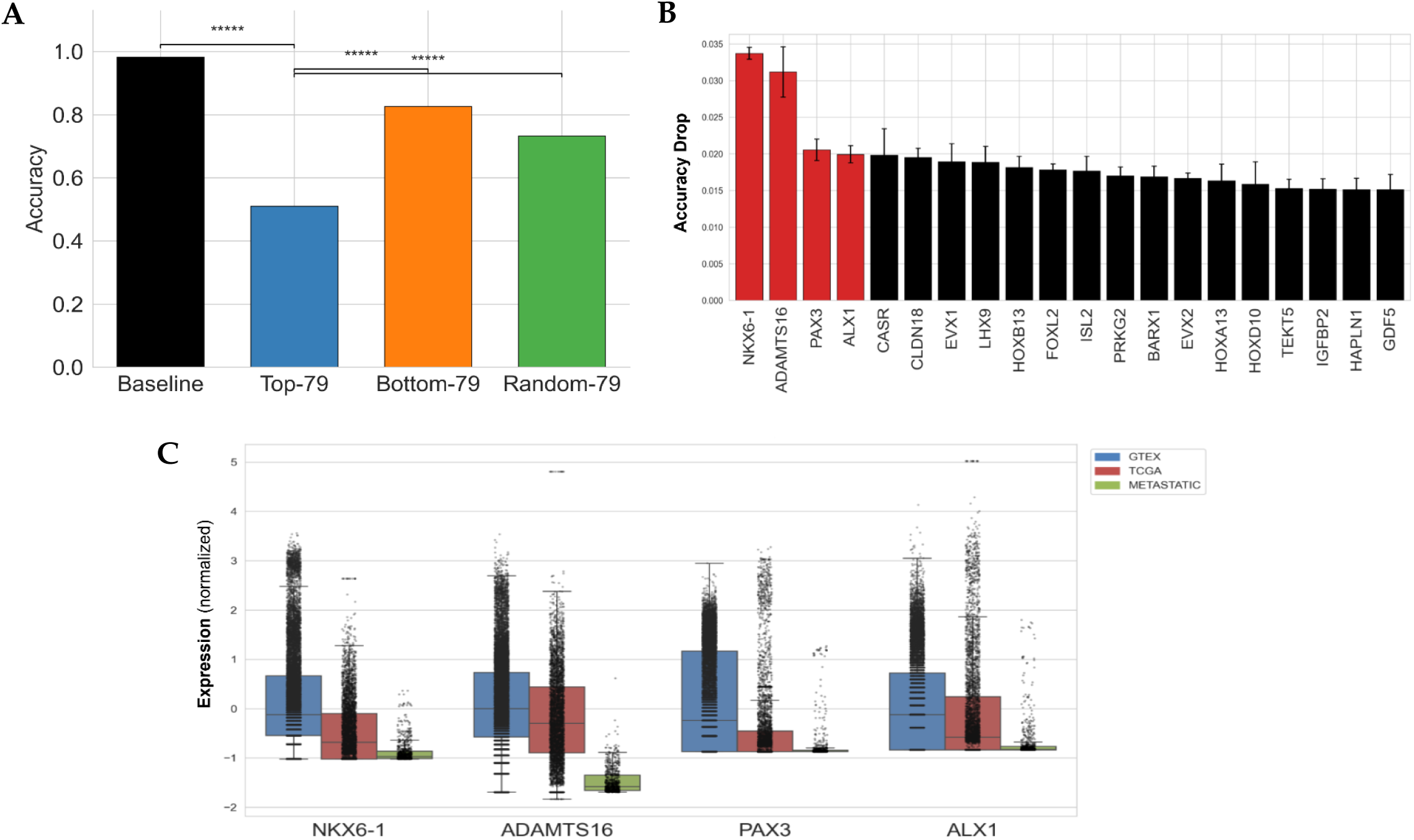
Monte Carlo Ablation. A – Bar plot of the most significant genes (n=79) affecting TCUP accuracy, (***** = p < 0.00001; McNemar’s χ² test). B – Top 20 most significant genes among them. 4 most significant highlighted in red C – Box plots of the top four genes’(red bars) expression (normalized, log-transformed) across all samples.

### 3.4 Primary and Metastatic BRCA

The ablation strategy was next employed to examine molecular distinctions between primary and metastatic breast carcinoma (BRCA). TCUP achieved high accuracy for both phenotypes (98.3% and 97.5%), making BRCA an ideal test case. Briefly, we iteratively pruned a gene set collectively to significantly impair metastatic BRCA prediction (p < 0.05, McNemar’s χ² test) while minimally affecting primary BRCA prediction (p > 0.05) (see Methods 2.6). This yielded a set of 19 genes (Fig. 5A). Literature review revealed that several of these genes have established tumor-suppressive roles in breast cancer dissemination, while others are less characterized (Fig. 5B). For example, the homeobox factors MSX1 and SOX30 are frequently downregulated or epigenetically silenced in metastatic tissue [36–38]. In contrast, some genes have been implicated in metastasis in other cancers but not in BRCA. SYTL1, a Rab27 effector involved in CD81⁺ exosome secretion and microenvironmental remodeling, has been shown to suppress lung adenocarcinoma spread [39] and correlate with improved survival in endometrial cancer [40], but its role in BRCA remains unreported. The identification of SYTL1—among a set largely composed of known BRCA-related genes—underscores the sensitivity of this ablation strategy in uncovering both established and previously unrecognized molecular contributors to metastatic biology.

**Fig. 5.**
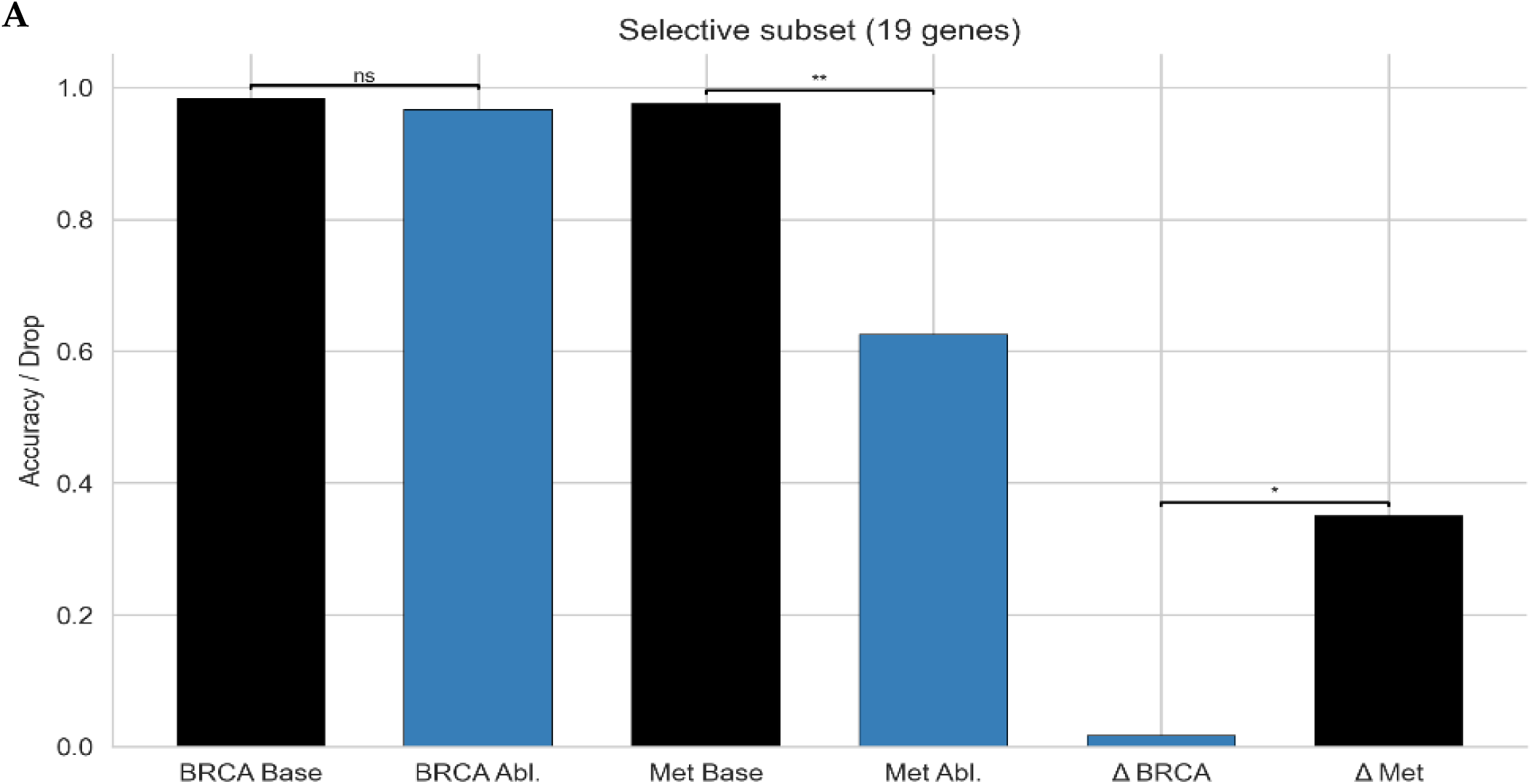
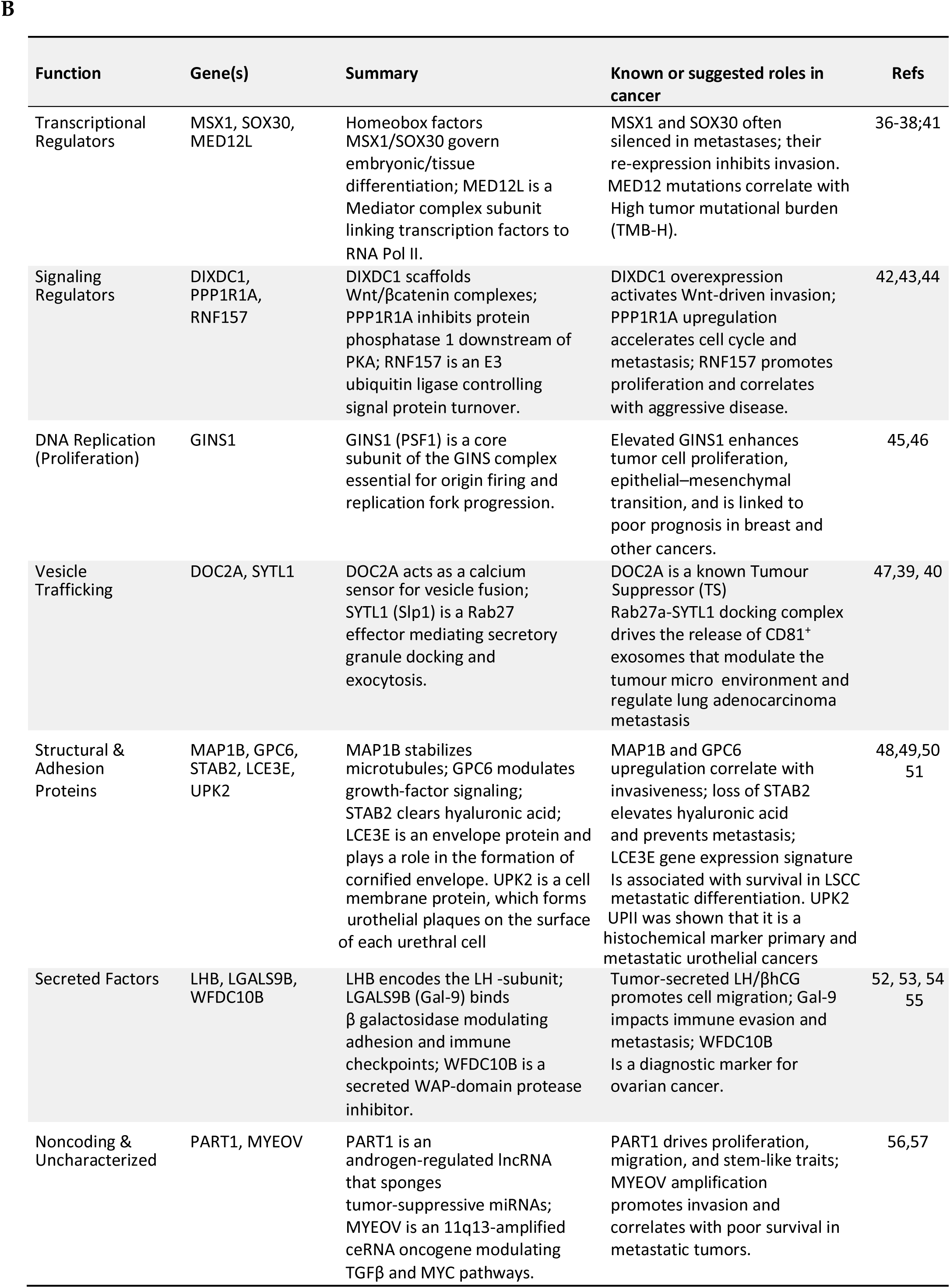
Differential Ablation. A – Differential ablation analysis between BRCA and Metastatic BRCA (Met) reveals 19 genes that significantly impact model performance for Met (*p < 0.05; McNemar’s χ² test; ** p<0.01; ns= non-significant, p>0.05), while exhibiting no significant effect on BRCA prediction accuracy (ns – non-significant). B – Summary table of selected genes, briefly highlighting their documented involvement in metastatic invasion and cancer prognosis.

### 3.5 Graphical User Interface

To facilitate external validation, we encapsulated the entire TCUP pipeline in a lightweight Dash web application (https://fohs.bgu.ac.il/rubinlab/TCUP/; source code available at GitHub, see Data and Code Availability). On the landing page (Fig 6A) users click Analyze to launch an overlay dialog (Fig 6B) where they drag-and-drop a transcriptome file—either a single sample or a matrix of multiple samples—and specify whether the cohort should be interpreted as Cancer (TCGA + metastatic reference) or Healthy (GTEx reference). Missing genes are automatically imputed with the appropriate cohort-median expression values to ensure robust prediction (Methods 2.5). After submission, the application renders a results panel (Fig 6C) that displays: (i) a bar chart of posterior probabilities across all tissue classes; (ii) the highest-probability label together with its absolute probability, TCUPs accuracy previously obtained for that label and a statement summarizing the number of genes imputed; and (iv) a ranked list of the twenty genes whose single-gene ablation most reduced classification accuracy for the predicted label, each annotated with a z-score reflecting how the input sample’s expression deviates from the gene’s standard deviation distribution across all samples (Methods 2.5), along with the corresponding p-value threshold. (* - p<0.05, ** - p<0.01, *** - p<0.001). In this way, the GUI delivers an immediately interpretable tissue-of-origin call while exposing candidate dysregulated genes that may guide downstream biological or clinical

**Fig 6.**
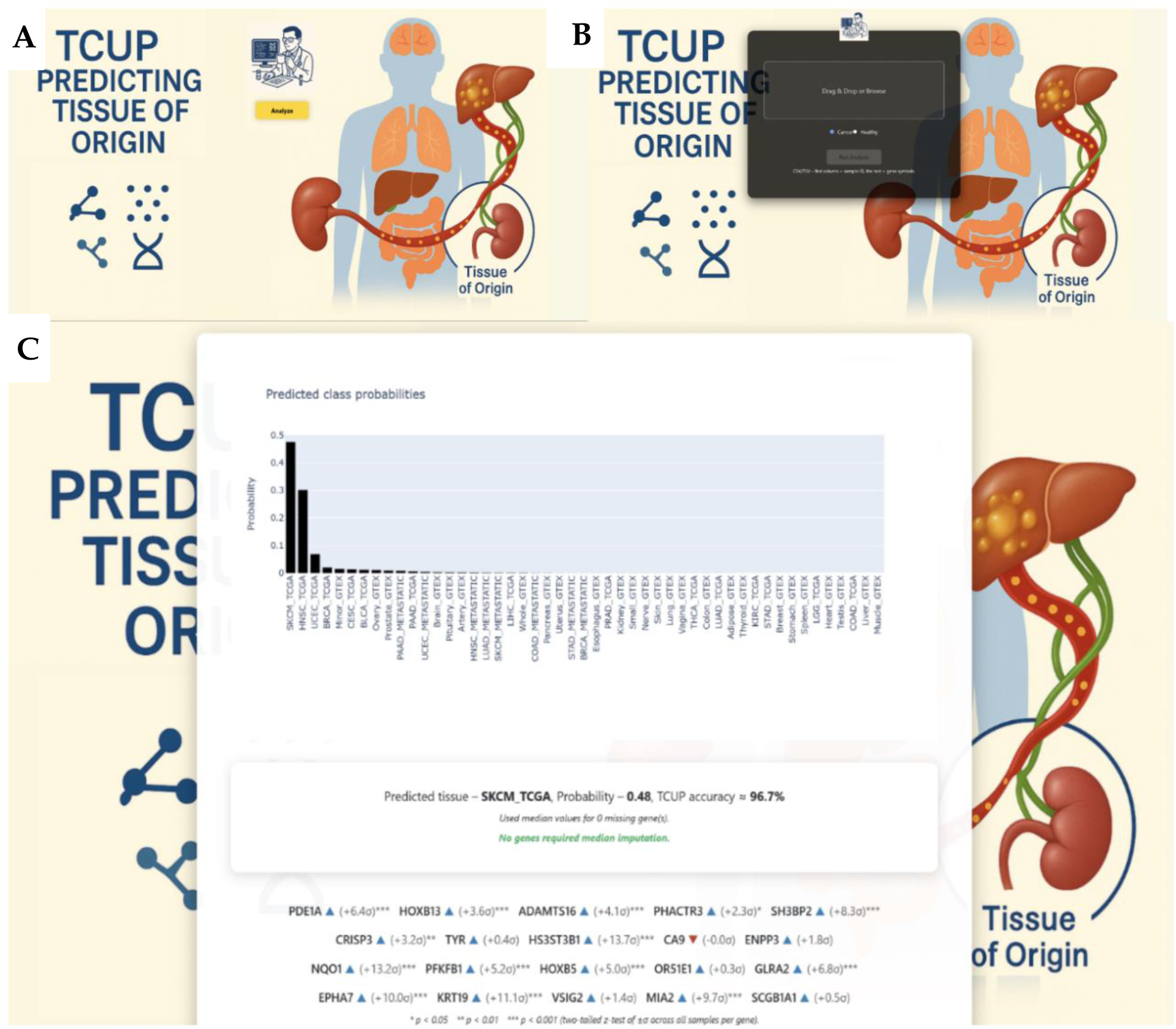
Graphical User Interface. A – TCUP landing page with an Analyze button that starts the prediction workflow. B – Overlay dialog in which the user uploads a CSV/TSV file (single or multiple samples) and selects the reference cohort (Cancer or Healthy). C – Example results page showing (i) a bar-plot of class probabilities, (ii) the predicted tissue with its probability and TCUP’s cross-validated accuracy for that class, and (iii) z-scored standard deviation of the input samples twenty genes whose single-gene ablation most reduced accuracy for the predicted label

## 4 Discussion

In this study we present a novel classification framework for TOO and CUP that combines Siamese neural networks with a contrastive auto-encoder and a meta-learning ensemble. Interpretability is achieved by Monte-Carlo gene-ablation analysis, which yields biologically and clinically meaningful insights. To ensure broad accessibility, we have released the resulting tool—TCUP— not only at the source code level but also as an open-access web application. Our results demonstrate that embedding the diverse transcriptomic features of healthy and tumour cells into compact signatures enables accurate discrimination across a broad spectrum of primary tumors and their advanced metastatic counterparts; the model’s strong performance underscores its utility in distinguishing primary from metastatic disease.

While TCUP matches or surpasses the performance of the most recent classifiers (Table 2), several limitations warrant consideration. Our “CUP” test cohort comprises metastatic biopsies obtained from anatomical sites distinct from their documented primaries. Although these secondary deposits are biologically more complex than resected primaries or locoregional metastases and thus serve as a useful proxy for validation, they are still not authentic CUP specimens. True CUPs display markedly higher genomic instability and inter-lesional heterogeneity than metastases of known origin [58], disparity thought to underline the persistent inability to identify their primaries. As a result, it remains to test the accuracy of TCUP with actual CUPs and can be expected to be lower than reported here. Moreover, because our inclusion criteria required metastatic biopsies taken from sites discordant with their documented primaries, the resulting “CUP” subset was comparatively small and unbalanced: only four metastatic tissue classes were available for analysis, totaling 120 samples— far fewer than the primary-tumour and healthy-tissue groups. Although we observed that CUP classes with larger sample counts (e.g., BRCA = 40, COAD = 35) tended to yield higher predictive performance, there is no assurance that the same relationship will hold for tumour types that remain under-represented and with low accuracy. In addition, the dataset is dominated by healthy and primary-tumour profiles, for which TCUP attains very high accuracies (GTEx = 99.3 %, TCGA = 96.2 %). Even with stratified evaluation, these disproportionately strong results can elevate aggregate metrics and may therefore overstate performance in the metastatic CUP context.

Many of the constraints noted above simultaneously confer important strengths. To mitigate the limited number of metastatic cases that met our inclusion criteria, we down-sampled over-represented TCGA classes, thereby preventing any single tumour transcriptome from dominating the latent space; this balancing step contributed to the comparably high macro F1-score achieved across primary tumors (96.2 %). In parallel, we broadened the transcriptomic spectrum by incorporating 26 anatomically distinct healthy tissues from GTEx, which improved separation within the embedded space and, to our knowledge, represents the first deliberate use of normal tissue to enhance CUP tissue-of-origin classifiers. Finally, accurate prediction is only the key; understanding the molecular basis of that prediction is the lock it must open before the door to rational CUP therapy can swing wide. By performing iterative and combinatorial gene ablations - including differential ablation comparing matched primary and metastatic labels - we embed a layer of interpretability within an otherwise complex ensemble, offering mechanistic insights that complement TCUP’s diagnostic utility. This analysis highlighted both established and emerging drivers of metastases, Well-documented regulators of metastatic progression—NKX6-1 [59, 60], ADAMTS16 [61], SOX30 [36–37] and MSX1 [38]—all surfaced among the top accuracy-affecting genes, each governing a distinct facet of tumour dissemination and proving critical to TCUP’s ability to differentiate tissues of origin. Their retrieval functions as an positive control, bolstering confidence in previously unrecognized candidates such as SYTL1, a Rab27 effector linked to lung-adenocarcinoma progression yet [36, 37], to our knowledge, not previously associated with metastatic breast cancer as we show here.

Future work will focus on addressing the limitations outlined above, beginning with the acquisition and integration of external CUP cohorts to further evaluate TCUP’s performance on these particularly heterogeneous tumors. We anticipate that routine use of the open-access portal (https://fohs.bgu.ac.il/rubinlab/TCUP/) will facilitate additional prospective validations as new cases are reported. Iterative retraining that incorporates nested CUP samples together with larger series of advanced metastases should improve latent-space separation and, consequently, classification accuracy. Finally, because multi-omics integration has usually been shown to outperform single-omics models [62, 63], we plan to develop a TCUP branch capable of ingesting alternative or combined data modalities, thereby expanding the tool’s applicability and—presumably—its predictive power.

## 5 Conclusion

TCUP shows that transcriptome-derived embeddings, paired with an interpretable ensemble architecture, can differentiate primary tumors from anatomically distant metastases while revealing gene-level signals that drive those distinctions. By rebalancing over-represented classes, adding normal-tissue profiles, and applying systematic ablation, the framework delivers both robust performance and biological context.

Released as an open-access portal, TCUP offers investigators an additional resource for CUP research and a foundation on which future studies may build toward biologically informed treatment selection.

## Code and Data Availability

All transcriptomic data used to develop TCUP were obtained from TCGA [30] and the curated training dataset is available upon request. TCUP is open-access and freely accessible at https://fohs.bgu.ac.il/rubinlab/TCUP/

## Supplementary

**Supplementary Figure S1.**
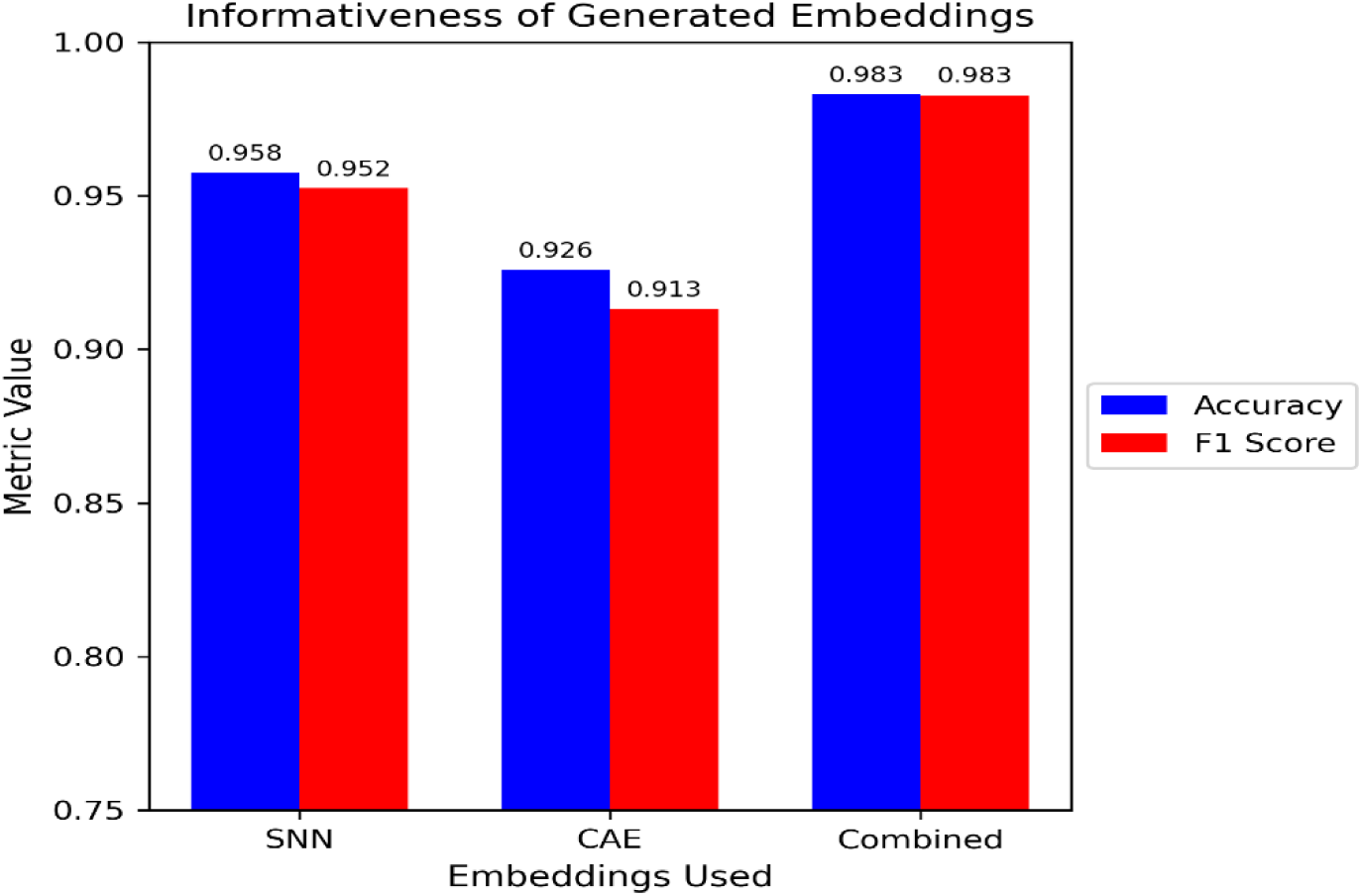
Informativeness of Each Embedding Set. Accuracy and F1 scores for network when using SNN alone, CAE alone or Combined embeddings

**Supplementary Figure S2.**
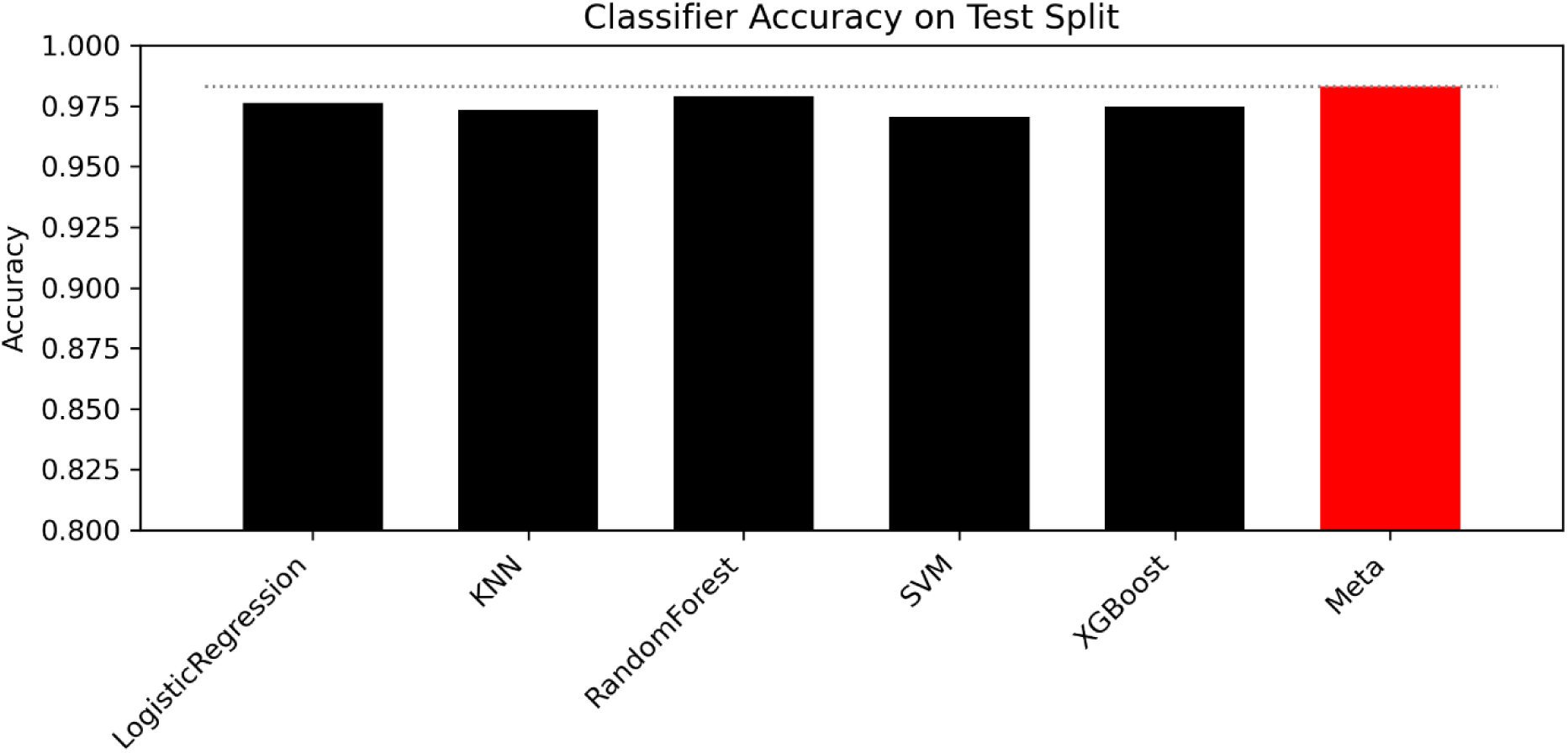
Meta Learner vs Standalone Classifiers. Accuracy of each classifier on combined embeddings alone and combined (meta learner) illustrating marginal yet informative increase in results.

**Supplementary Figure S3.**
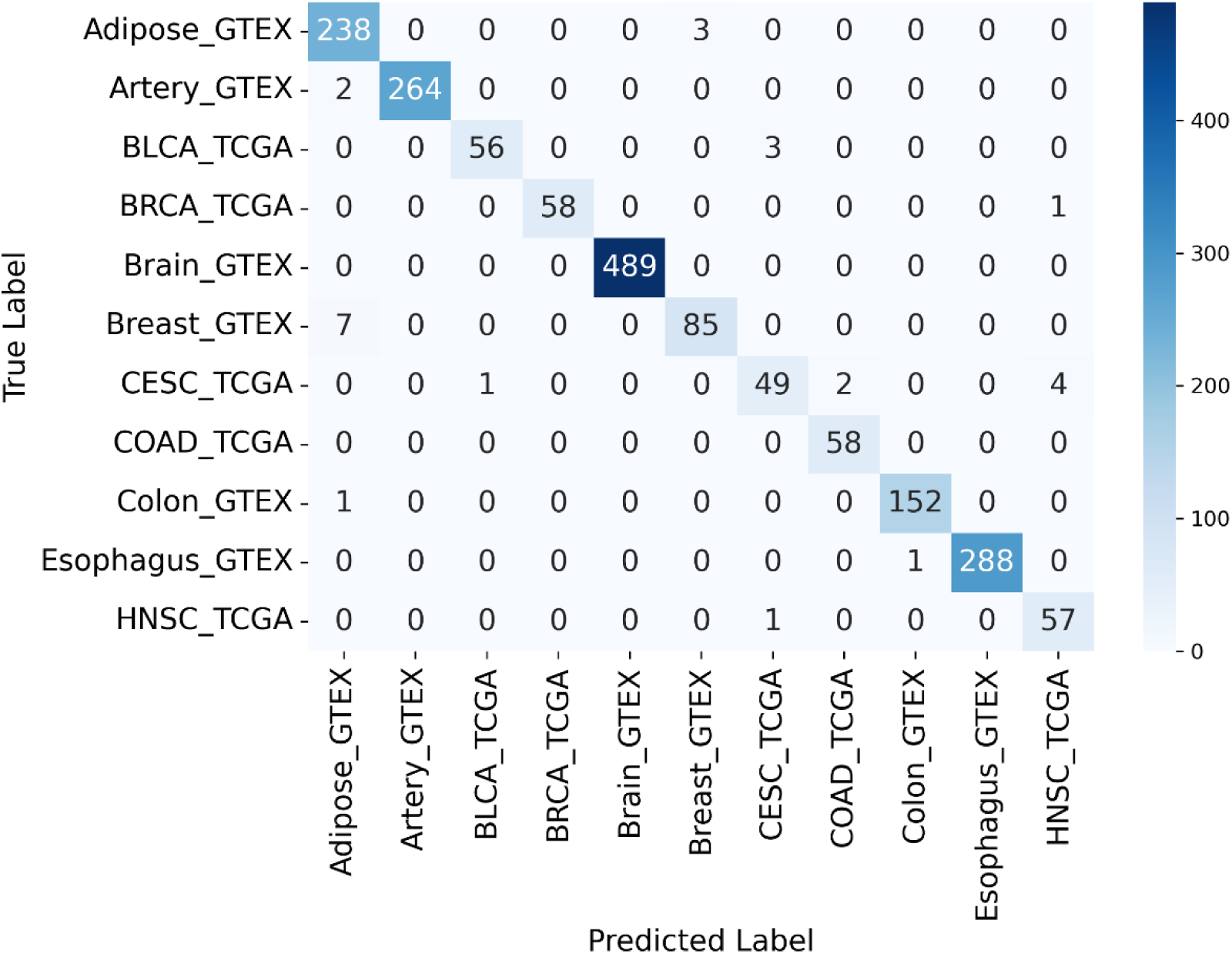
zoomed in confusion matrix of figure 3A (1-11)

**Supplementary Figure S4.**
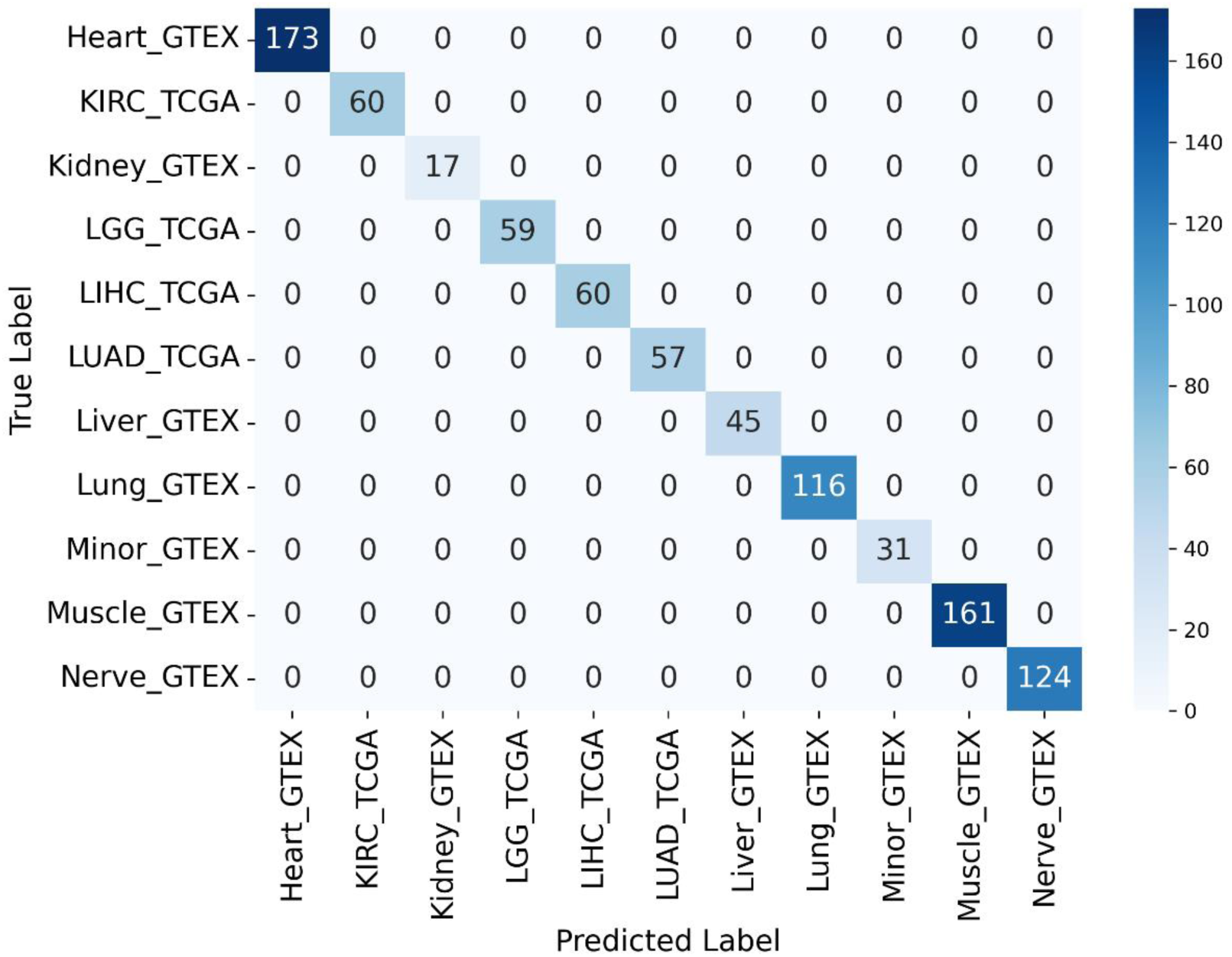
zoomed in confusion matrix of figure 3A (2-22)

**Supplementary Figure S6.**
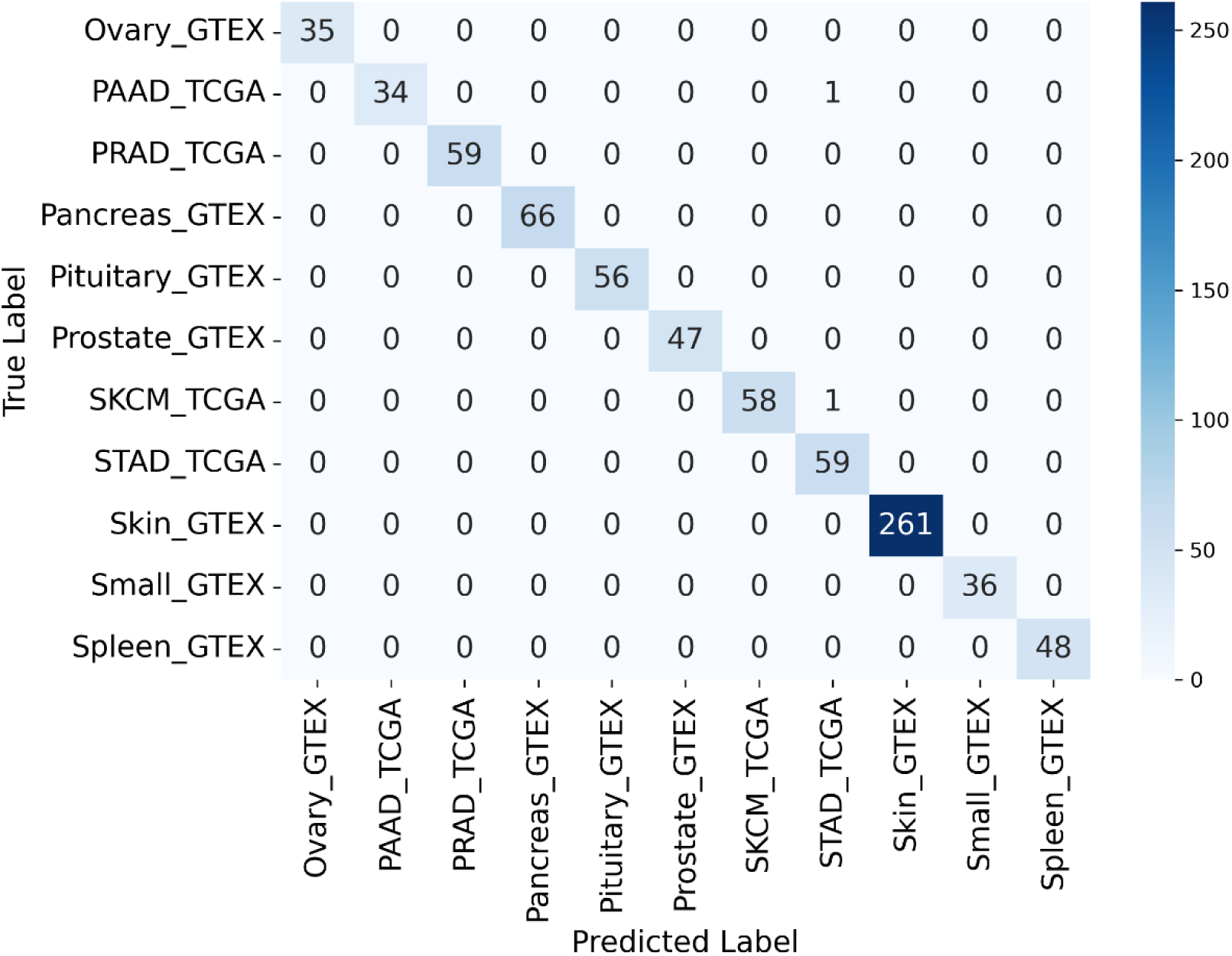
zoomed in confusion matrix of figure 3A (3-33)

**Supplementary Figure S7.**
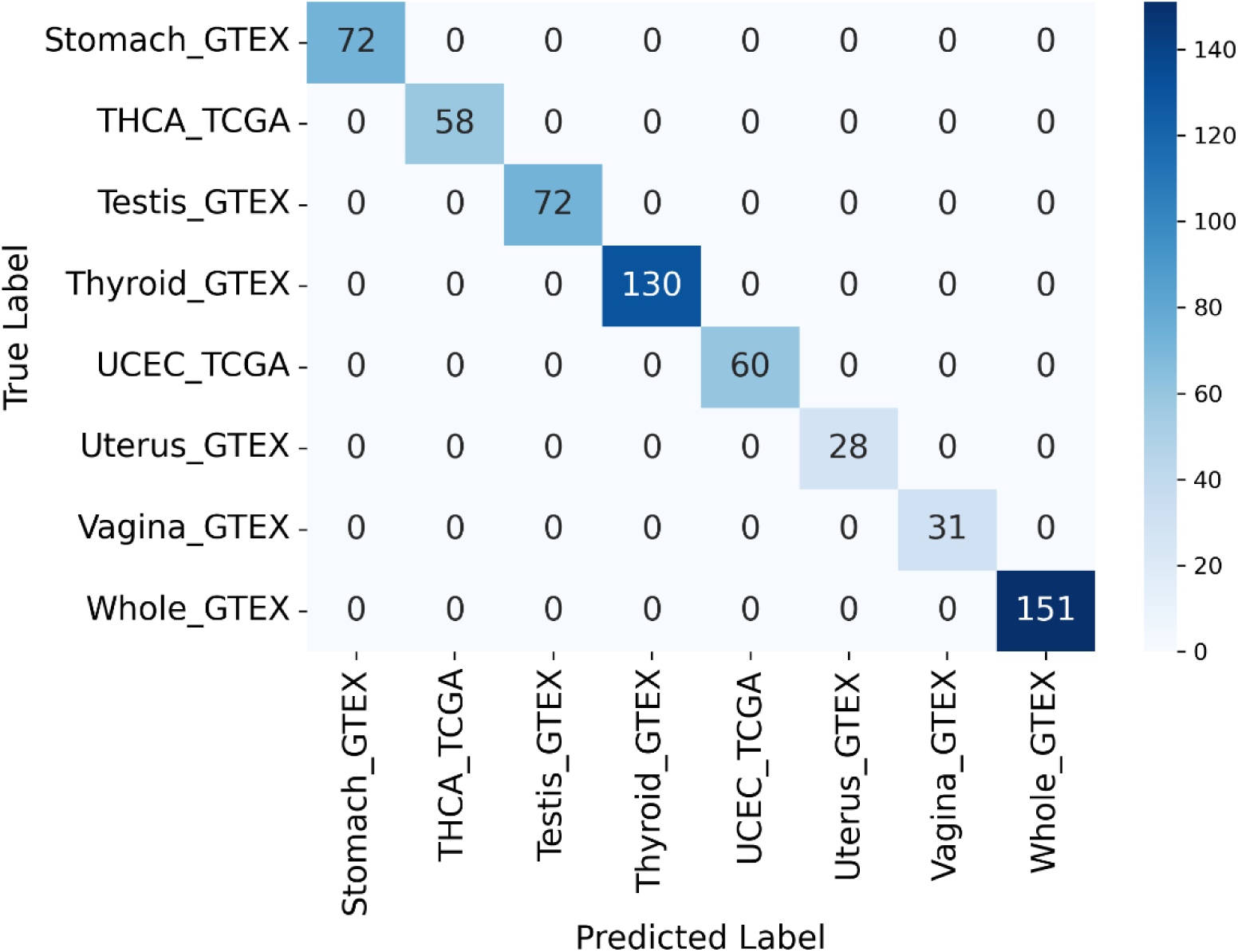
zoomed in confusion matrix of figure 3A (34-41)

**Supplementary Figure S8.**
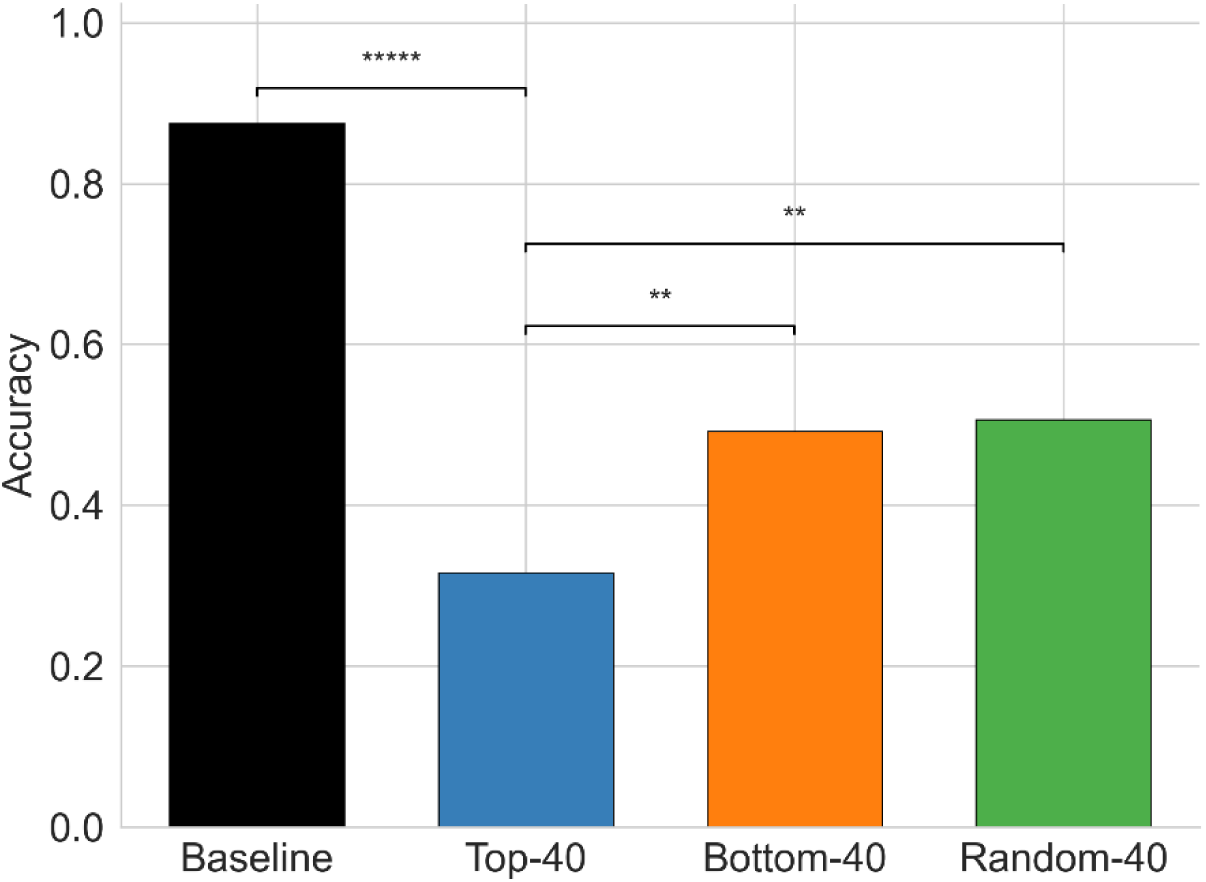
Most significant gene set affecting TCUPs metastatic accuracy.

**Supplementary Figure S9.**
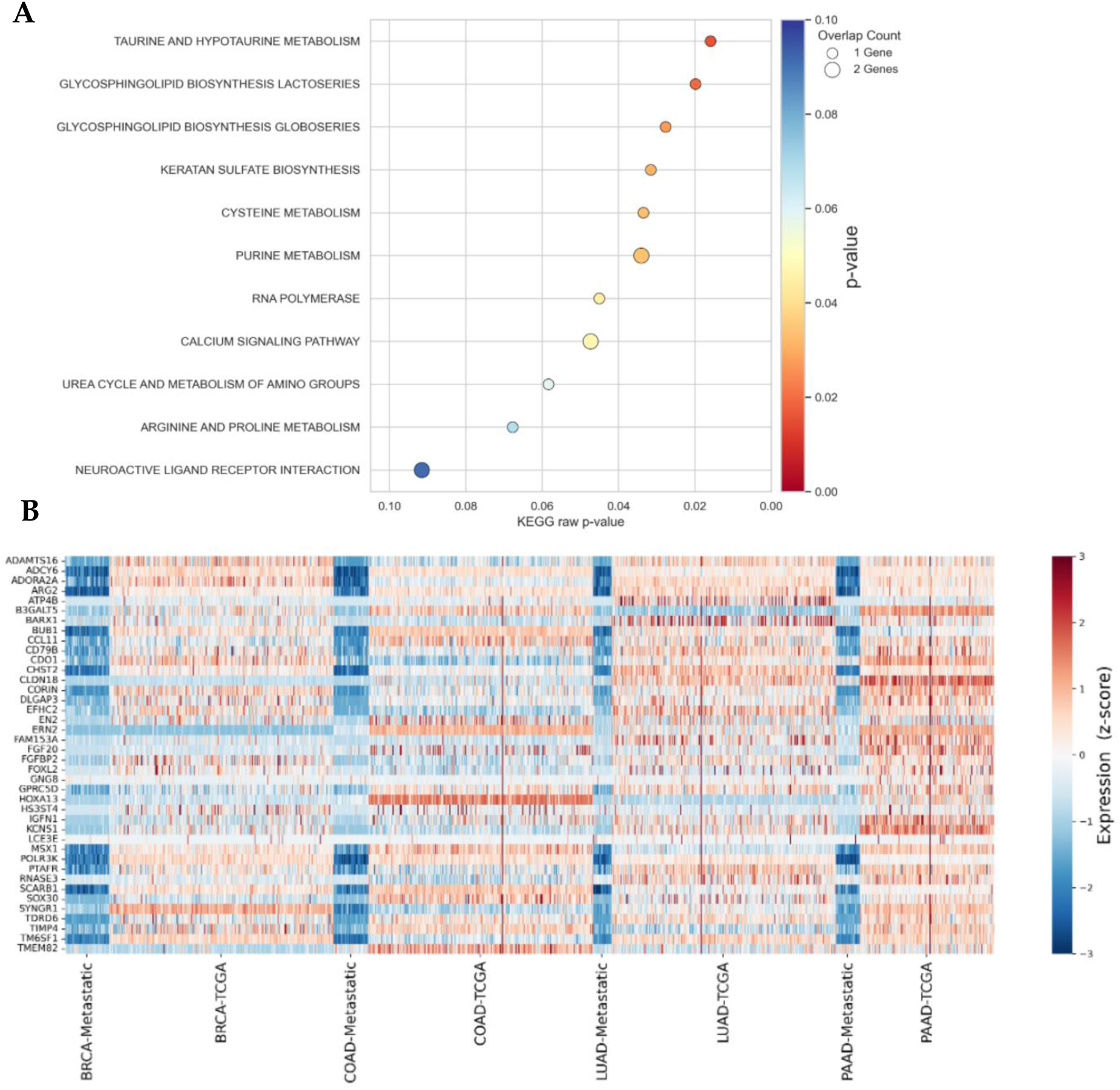
Functional enrichment of the metastasis-associated gene set. A. Bubble plot of KEGG pathways enriched among the top metastatic genes (n = 40); circle size reflects the number of genes per pathway and color encodes raw p-value (threshold p < 0.10). B. Heat-map of the 40 genes across primary and metastatic labels (BRCA, COAD, LUAD, PAAD); values are z-scores relative to the cohort-specific (samples in the heatmap) median expression.

